# Whole-body coarticulation reflects expertise in sport climbing

**DOI:** 10.1101/2025.01.10.632403

**Authors:** Antonella Maselli, Lisa Musculus, Riccardo Moretti, Andrea d’Avella, Markus Raab, Giovanni Pezzulo

**Affiliations:** Institute of Cognitive Sciences and Technologies, National Research Council, Rome, Italy; Department of Performance Psychology, Institute of Psychology, German Sport University Cologne, Cologne, Germany; Department of Biology, University of Rome Tor Vergata, Rome, Italy; Santa Lucia Foundation, IRRCS, Rome, Italy

**Keywords:** coarticulation, whole-body kinematics, complex motor skill, climbing, planning

## Abstract

Taking sport climbing as a testbed, we explored coarticulation in naturalistic motor-behavior at the level of whole-body kinematics. Participants were instructed to execute a series of climbing routes, each composed of two initial foot-moves equal in all routes, and two subsequent hand-moves differing across routes in a set of eight possible configurations. The goal was assessing whether climbers modulate the execution of a given move depending on which moves come next in the plan. Coarticulation was assessed by training a set of classifiers and estimating how well the whole-body (or single-joint) kinematics during a given stage of the climbing execution could predict its future unfolding. Results showed that most participants engage in coarticulation, with temporal and bodily patterns that depend on expertise. Non-climbers tend to prepare the next-to-come move right before its onset and only after the end of the previous move. Rather, expert-climbers (and to a smaller extent, beginner-climbers) show early coarticulation during the execution of the previous move and engage in adjustments that involve the coordination of a larger number of joints across the body. These results demonstrate coarticulation effects in whole-body naturalistic motor behavior and as a function of expertise. Furthermore, the enhanced coarticulation found in expert-climbers provides hints for experts engaging in more refined mental processes converting abstract instructions (e.g., move the right hand to a given location) into motor simulations involving whole-body coordination. Overall, these results contribute to advancing our current knowledge of the rich interplay between cognition and motor control.

**NEW & NOTEWORTHY:** The current study explores the way in which having formed a plan for a sequential motor task affects its execution. We showed that climbing expertise increases the extent to which participants adjust their motor execution based on the moves that follow in a planned route. These results provide evidence of coarticulation in naturalistic motor behavior and suggest enhanced skills in mentalizing forward motor control and optimal-control strategies in expert climbers.

## INTRODUCTION

Naturalistic motor behavior is strictly intertwined with the cognitive processing underlying decision-making and planning. During everyday activities (such as walking or driving through a busy road) as well as when executing skilled actions (like climbing on a rocky wall), motor control, planning, and decision-making processes are deeply entangled (1–4). In all cases, a successful execution requires the transformation of an abstract plan into an ordered sequence of bodily actions. For example, when expert climbers are faced with a route they plan (i) the sequence of holds to be grasped, (ii) which limbs should move one hold to the next, and (iii) how to coordinate the whole body actions to optimally execute the planned sequence of actions (5,6). On the other hand, the actual execution of naturalistic actions is subject to some degree of intrinsic motor variability (7) that makes the exact dynamics of the bodily state (e.g., its posture and dynamics, fatigue and psychological attitude) unpredictable, at least to some extent. Such a degree of uncertainty implies the need for online monitoring and may require online updates of the initial plan (8,9,6). The study of how planning, decision-making and motor control interact reciprocally is crucial to gain a comprehensive understanding of human behavior and to move beyond the classic approach of testing each aspects in isolation under highly controlled setups (10). The current study aims at providing a contribution in this direction.

We still have a limited understanding of the complex interplay between motor control, planning and decision-making in naturalistic behaviour. Classic decision-making models account for how decisions are made under uncertainty, with either a serial or parallel view on the decision process (11). These models, however, do not account for the impact of the constant stream of sensorimotor information gathered during ongoing actions (12–16). Recent approaches have started describing the link between motor control and planning as a continuing, dynamically evolving, and constant feedback loop between sensorimotor and cognitive input (5,17,18). The goal is to elucidate how bodily constraints, (past) sensorimotor experience, as well as motor skills interfere with information gathering and affect cognitive planning and the decisions made at each step of naturalistic action sequences (19). Better understanding the interplay of motor and cognitive processes is crucial for understanding how humans act in the real world (2), and how they develop their cognitive and motor prerequisites for (6) and become experts in (20) complex actions.

Researchers in motor control started investigating how movements and decisions interact with each other (21). It was shown how cognitive states – e.g., uncertainty, motivation and decision’s confidence – affect the motor output of a given choice, in particular movement trajectory, speed and vigor (22–25). Also the opposite line of cause-effect has been investigated: the motor state of the agent and the related perceived effort were shown to have a significant impact on online decisions, leading in some cases to changes of minds, e.g., when deciding which of two targets to reach or which trajectory strategy to pursue (26–29). Most of these studies worked on controlled laboratory setups and focused on reaching tasks. Many other aspects of the interplay between planning, decision-making and motor control remain to be explored. Here we move a step forward in this direction by focusing on a complex (whole-body) naturalistic motor task, specifically testing for the impact of a full planned sequence of actions on the execution of intermediate steps. For this, we take sport climbing as a representative instance of naturalistic embodied sequential task in which individual expertise is expected to play a key role.

A well-known signature of the impact of planning on motor behavior can be found in the phenomenon of coarticulation (30). A long tradition has identified and studied coarticulation in the context of speech production (31,32). The phenomenon consists in the modulation of the way a given phoneme is pronounced depending on what follows next in a word. More generally, coarticulation refers to a phenomenon in which the execution of subtasks is influenced by the overall task (33). Beyond the case of speech production, coarticulation has been reported for a variety of sequential tasks retaining a link to language processing – e.g., fingerspelling (34) and sign language (35) – or not – e.g., finger-tapping, keys-pressing (36) and reaching sequences (37,38). In all cases, it was found that the execution of a first common part of the sequence is adapted according to what follows in the sequence. For example, Kalidindi and Crevecouer (2024) showed that during a sequence of two target reaching task, both the speed profile and the path of the trajectory reaching to the first (common) target are influenced by the location of the second target, in a way that can be predicted by an optimal feedback control model. The key point is that coarticulation in any form provides implicit evidence for the agent having a plan representation that encompasses the future unfolding of sequenced task, and is thus a valid tool for exploring the interplay between planning and motor control.

Coarticulation effects have been reported in relatively simple motor tasks that do not involve whole body coordination (e.g. finger tapping, key pressing, and reaching sequences). In this study we took sport climbing as a testbed for studying coarticulation in whole-body naturalistic motor behavior. Sport climbing provides a two-fold advantage. First, it offers a window into a complex motor skill that is intrinsically sequenced: each end-effector (hands or feet) must move from a position to the next, and typically (although not always) end-effectors move one at the time in serially sequenced patterns. Second, as in other sports, expertise and task difficulty can be well characterized (39,40). Specifically, climbing expertise encompasses physical ability in performing challenging whole-body movements, as well as perceptual and cognitive abilities in reading out the possibilities of action afforded by a climbing wall (41,42), in creating climbing plans that get progressively more sophisticated with learning (5,43), and in memorizing routes (44).

We aim to test two hypotheses. The first hypothesis is that once climbers have formed a plan for how to execute a route – e.g., when explicitly instructed about the sequence of movements to be executed – they will show signatures of coarticulation in the route execution. The second hypothesis is that the degree of coarticulation is modulated by the climber’s expertise, with expert climbers engaging in a higher degree of coarticulation due to their enhanced ability in forming refined plans involving motor simulations of the climbing route execution at the level of the whole body (5,18,44,45).

To address whether and how climbers with different degrees of expertise engage in coarticulation when executing a planned climbing route, we designed a task that could be feasible for participants with no climbing experience and still complex enough to allow coarticulation to emerge. Participants had to execute a series of climbing routes (80 trials including 10 repetitions for 8 different routes). All routes consisted in a common sequence of two initial foot-moves, followed by two hand-moves differing in each condition according to the combination of the hand that had to perform each move and the location of the second (final) hold to be grasped (Figure 1). While opting for more complex and challenging routes would have been ideal for enhancing the amount of motor coarticulation, the choice of simpler routes was necessary to allow non-climbers to successfully execute them – and hence to test for the impact on expertise on the degree of coarticulation.

**Figure 1.**
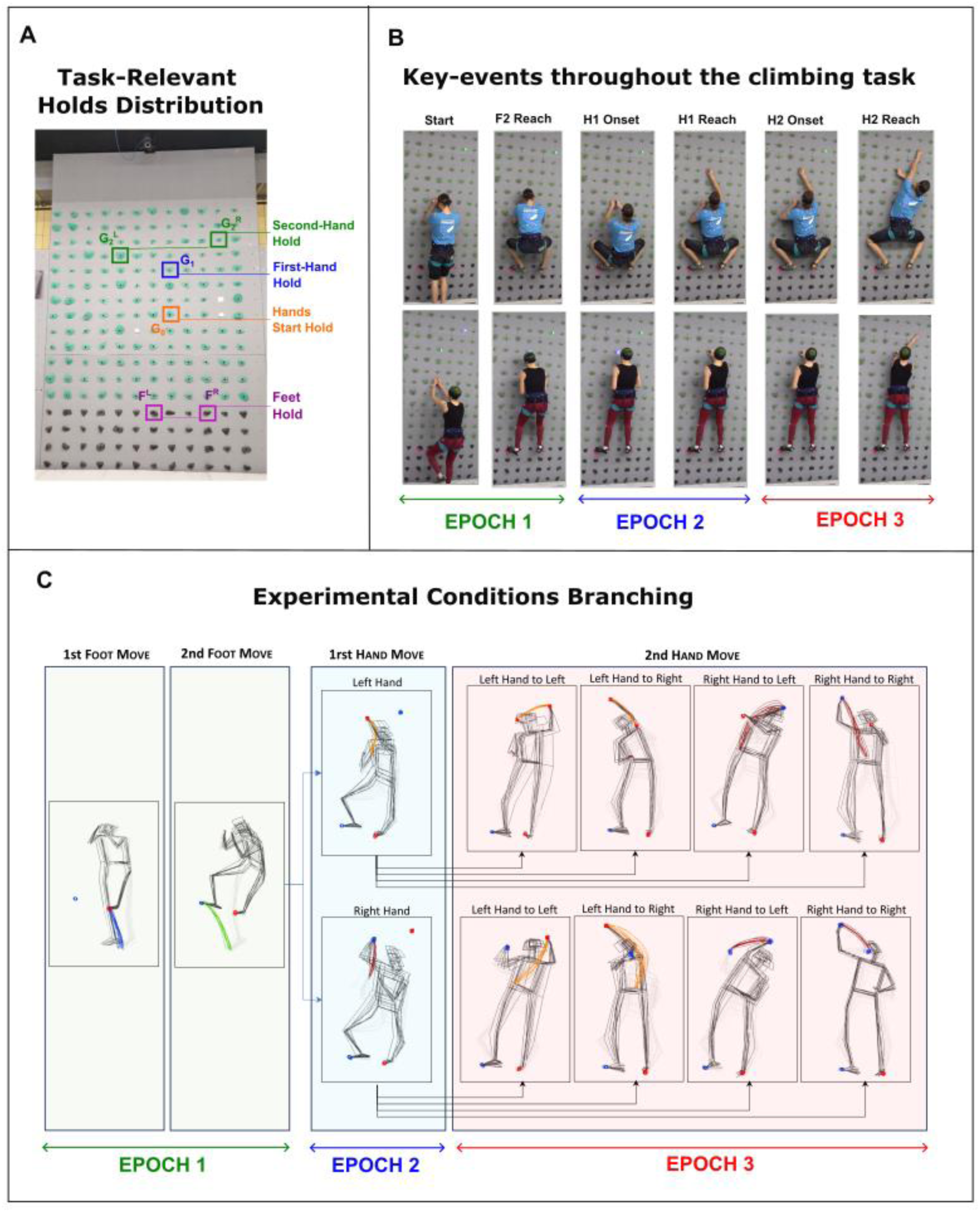
Climbing task. (A) Climbing wall utilized in the experiment; the holds available for the routes are highlighted by the coloured squares: the holds to be reached in the two foot-moves (F^L^, F^R^), the hold to be grasped with both hand in the starting posture (G_0_), the hold to be reach in the first hand-move (G_1_), and the two possible hold targets of the second hand-move (G^L^_2_, G^R^_2_). (B) Snapshots of the climbing execution at key-events for two representative participants, an *expert-climber* (upper row) and a *non-climber* (lower row); for space reasons the key-events associated with first-foot reach and second-foot onset are omitted. (C) Overall view of how the 8 different experimental conditions branch out.

The degree of coarticulation was assessed by analyzing the whole-body kinematics (reconstructed from the optical tracking of 25 markers placed throughout the body) recorded during each trial execution. Differently from previous studies that focused on pre-selected and/or summary kinematics variables such as the center of mass (COM) (43,46) or specific joints trajectories (47), we aimed at assessing the richer behavior expressed in whole-body coordinated movements, which may bring along a wider and finer set of information for inspecting the role of motor skill, and of inter- and intra-individual motor variability. The whole-body kinematics across different temporal intervals was analyzed to assess how accurately and which aspects of the kinematics up to a given stage of the climbing execution (e.g. before the onset of the first hand-move) allowed to predict the following unfolding of the route (e.g. which hand will perform the first hand-move). To test for the role of expertise we recruited 21 participants with different climbing experience, ranging from non-climbers to expert-climbers, and grouping them in three categories: *non-climbers*, *beginner-climbers*, and *expert-climbers*.

Looking for coarticulation in the whole-body kinematics implies having to deal with high dimensional data. We addressed this issue by deriving low-dimensional descriptions of the whole-body kinematics obtained with spatiotemporal principal component analysis (stPCA) (48,49), and use these compact descriptions to train a set of linear discriminant classifiers that provide quantitative estimates of the extent and the way in which planning-ahead for future moves affects the kinematics during the climbing execution.

Overall, our study provides a direct test of the impact of planning on naturalistic whole-body motor behavior, and novel evidence for how planning induces coarticulation in the kinematics of climbing tasks execution in a way that is considerably modulated by the climber’s expertise.

## MATERIALS AND METHODS

### Experimental design

Participants were asked to execute instructed climbing routes on a vertical instrumented laboratory-based wall. The tasks included 80 trials (10 repetitions of 8 different routes). The spatial configuration of the holds is shown in Figure 1. For all trials, the sequence started by standing in front of the wall grasping with both hand a hold placed 120 cm above the floor level (G_1_). Next, participants had to move both feet on two dedicated holds (60 cm high), and continue with the two hand-moves. The first hand-move, performed either with the right or the left hand, always reached the same target hold (G_1_), 180 cm high. The target of second hand-move, instead, could be one of two holds: G_2_^L^ located 80 cm left and 20 cm above G_1_, and G_2_^R^ located 40 cm to the right and 40 cm above G_1_. This configuration was chosen to have routes feasible for participants of different expertise levels, from novices to experts.

The experiment had a 2 x 2 x 2 design, with the three factors being: *H1-Hand*, the identity (right or left) of the hand performing the first hand-move; *H2-Hand*, the identity of the hand performing the second hand-move; *H2-Target*, the final hold target. All factors have two levels: *right* vs *left*. Participants performed 10 repetitions of each condition for a total of 80 trials. A complete trial consists in a first part including the two foot-moves, followed by the first hand-move executed with the right (left) hand in half of the trials. For each of these two subsets, the third move could be one of the four possible combinations of *H2-Hand* and *H2-Target*. Trials were presented in a pseudo random order, with a randomly ordered presentation of the 8 conditions included in our experimental design within each of the 10 blocks of trials. The experimental protocol was submitted to, and approved by, the local university’s ethics committee (no. 022/2021).

### Experimental setup and procedure

The experimental setup consisted in an infrared camera based VICON system for motion tracking including 10 infrared cameras (119.88 Hz, VICONTM, Oxford, UK), integrated and synchronised with a sensorised climbing wall (2.40–3.60 m; ClimbLing) equipped with touch-sensitive, lightning up LEDs. The ClimbLing wall allows administering a given climbing problem by lightning up (any) sequence of holds, colour-coded according to the hand that should grasp them (blue for right, green for left). The motion tracking and wall systems run on two dedicated PCs. The setup further included a tablet used to administer the demographic form and to enter the body metrics (weight, height, ape index and grip force) measured by the experimenter.

The procedure started with instructing the participant about the climbing task, and having them to sign the consent form, and to fill a demographic questionnaire including the following information: age, type and frequence of sport practice, and experience with sport climbing. Next, the participant was equipped with a set of 25 reflective markers distributed throughout the body with a spatial configuration optimally costumed so to mitigate the problems associated with the occlusion of the front body from the wall itself. Before starting the climbing session, anthropometric measurements including high, weight, and arm span were collected with measurements of the grip force for both hands.

In all trials participants started facing the wall at about 1 m away from it, holding a T-pose. The climbing problem was shown lighting the holds accordingly. Upon the experimenter giving a GO signal, participants had to execute a *mimicking* of the climbing hand-moves while staying at place. Next, they were instructed to move the hands down, walk towards the wall and reach the starting position with both hands on the starting hold (G_0_) and both feet on the floor and close by the wall. Next, they were instructed to start the climbing execution and complete it at their own pace.

The *mimicking* phase was included with a twofold purpose. First, to ensure that participants formed a clear climbing plan before starting the actual execution. Second, to explore possible correlations between mimicking, a common practice among expert climbers, and actual execution performance. This analysis, as well as the analysis correlating performance and idiosyncrasies in climbing strategies with body metrics and practice experience, is left for follow-up studies.

### Participants

21 participants (6 female; age 25.8 ± 3.6 years; height 177 ± 8.9 cm; weight 71,0 ± 11,9 Kg), all right-handed, were recruited by posting a call at the local university campus and at a nearby climbing gym. One participant did not finalize the experimental session, performing only 4 of the 10 blocks, because of the task being too challenging to him; his data were excluded from the analysis. Another participant completed only 8 out of the 10 for time constraints and his data were included in the analysis.

Participants were initially recruited as *beginners* and *experts*. Expertise was assigned based on the following criteria: experts should have 5+ years of practice in climbing or bouldering and be able to climb routes with difficulty 7a (in the French scale) or higher. Beginners should have 1 year or less of practice and be able to climb routes with difficulty no higher than 6a. Still so, the recruited participants were not segregated into two clearly separated groups of experts and beginners but they were rather continuously distributed in terms of expertise. We therefore grouped them into three re-defined expertise groups: *non-climbers*, who never practiced climbing; e*xpert-climbers*, who practiced climbing since at least four years and can perform routes of difficulty 7a+ or higher; *beginner-climbers*, all others.

### Data collection and pre-processing

For each participant data collection included the demographic, body metric and sport related information recorded before the experiment’s start, and the whole-body kinematics collected during the experiment. The kinematics of the 25 markers, placed on key joint locations throughout the body, was recorded with a set of 10 VICON infrared cameras system at 120Hz and with an average spatial precision below 3 mm.

The kinematics recordings were first pre-processed with the Vicon Nexus software. An ad-hoc whole-body model identical to all participants was used as a template to reconstruct and pre-label the markers. Next, each trial was visually inspected to check and correct the automatic labelling (i.e., fill the missing labelling and fix the mislabelled markers). A gap filling procedure was then run, and data were exported into c3d format.

### Data analysis

The kinematics data were analysed with a customized Matlab® pipeline. Kinematics data were band-passed filtered with a zero-lag Butterworth filter of order 3 and lowpass frequency 10 Hz (Matlab *butter* and *filtfilt* functions), as done in standard biomechanical procedures of kinematic data filtering to remove recording noise and to guarantee smoothness for differentiation (50). Positional data were next differentiated (Matlab *diff* function) to get velocities and accelerations. The kinematics recorded throughout each single trial was split in two parts: the mimicking and execution phases.

Trials in which participants did not correctly follow the problem instructions (e.g., swapping the hands order or moving to the wrong final target) were removed from the analysis. A total of 32 trials were removed from the statistical analysis, corresponding to about 2% of all the trials recorded.

### Segmentation of the climbing execution

The recorded climbing execution was segmented based on its four sub-movements: first foot-move (F1); second foot-move (F2); first hand-move (H1); second hand-move (H2). The four moves were all delimited on the lower end by the onset of end-effector’s movement, and on the upper end by its arrival at target. In addition, a climbing *start* event was identified as the time in which both feet were on the floor and close to the wall, both hands were holding the initial hold *Hold_0* and hands’ and feet’s speeds were below 0.15 m/s. In the ideal condition in which the participant complied with instructions, the *start* event corresponds to the onset of the first foot-move. In some cases, however, participants did not comply with the experimenter instructions and started the climbing directly from walking, i.e., skipping the initial posture. In these cases, the *start* event was identified as the time in which the heel of the stepping foot gets at 0.5 cm from the wall, while the remaining three end-effectors comply with the standard constraints on position and speed. Figure 1B shows six out of this eight delimiter stages for two representative participants. The *start* position depicted in the lower row provides an example of trial in which the participant did not comply with the instructions.

The segmentation was performed via an automated and customized Matlab® pipeline. Key-events were identified based on thresholds constraints on the speed of the end-effector and its position relative to the relevant landmarks (e.g. a specific hold location, the wall, or the floor). The results of the automated segmentation were visually inspected to correct for possible misidentifications. The visual inspection consisted in examining visualizations of the body posture and the end-effector trajectories in between them for each key-events, and was done for each of the 8 conditions and for each participant (an example of the visualizations used is shown in Figure 2). This procedure further served to verify that participants complied with instruction executing the task as a sequence of serial end-effectors moves.

**Figure 2.**
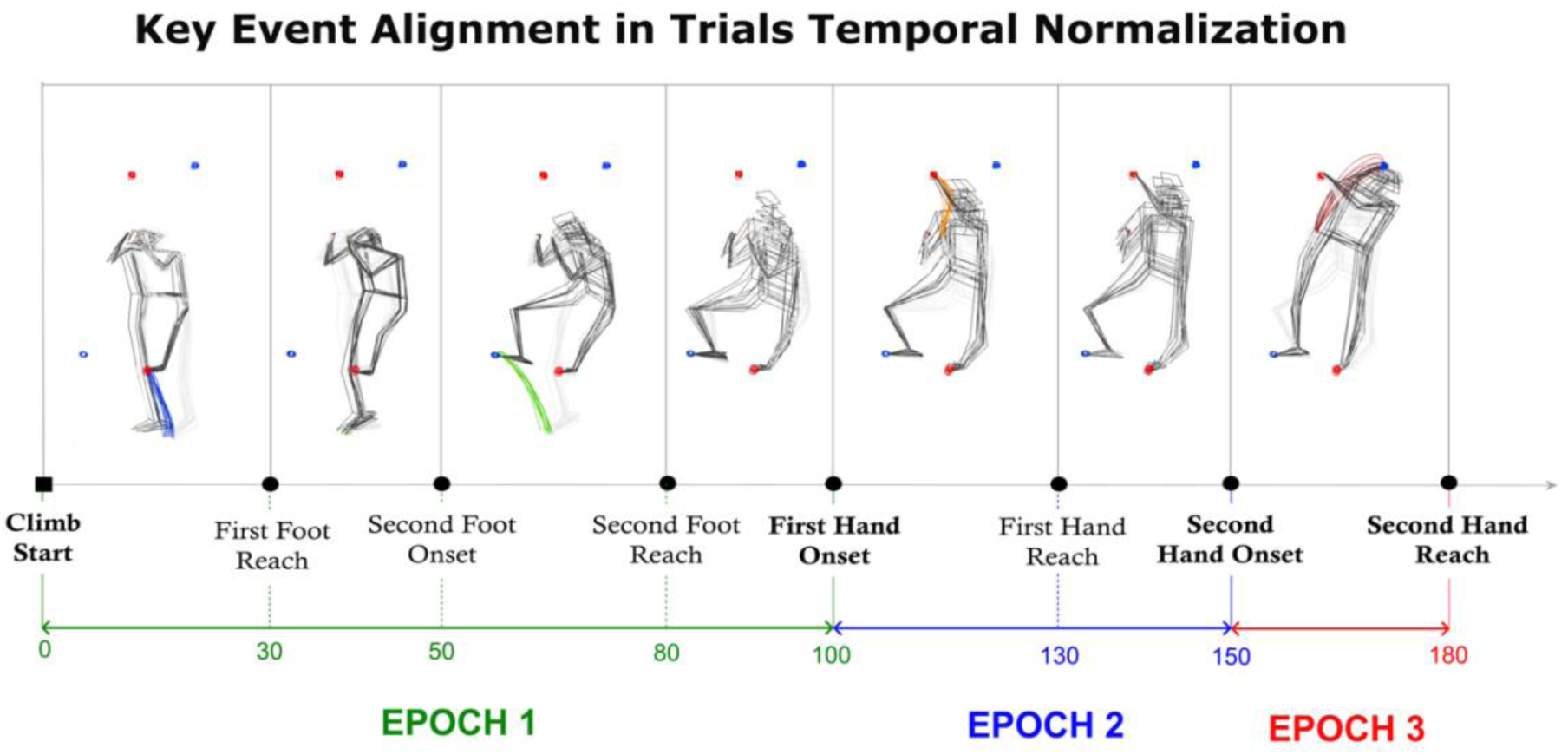
Alignment of key-events. The figure shows the posture of a representative participant at the key-events of the climbing task execution as extracted from motion capture data. For all participants, each trial from the start (corresponding to Climb Start, i.e. the onset of the first foot-move) to the end (corresponding to the reaching of the second hand-move) of the climbing execution was resampled on 180 time frames, and each key-event was aligned resampling the intervals of end effectors’ moves (from onset to reach) onto 30 time frames, and the adjustment phases (from the reaching of an end-effector’s move to the onset of the next move) onto 20 time frames.

### Assessments of coarticulation

The whole-body kinematics data were initially analysed at the individual level to derive individual indexes of the degree of coarticulation. Analysis at the group level was done in a second stage to performed statistical comparisons of individual indexes across participants and experimental conditions.

#### Individual coarticulation index

To quantify the degree of coarticulation we performed a linear discriminant analysis (LDA) (51). By definition, LDA quantifies the accuracy with which it is possible to assign an observation to a group (or class) given its properties. In running the LDA, we take as observations all the trials from a given participant, characterize them with the whole-body kinematics recorded in a given time-interval across the climbing execution, and use them to predict the type of moves that will follow. For example, taking as observation the whole-body kinematics from the climb’s start to the onset of the first hand-move, we could train a LDA classifier to predict which hand will perform the first hand-move.

Different linear discriminant classifiers were trained with the whole-body kinematics spanning different temporal intervals, as well as with subsets of single-joints kinematics. Three classification problems were tested independently: *H1-hand*, the identity of the hand performing the first hand-move; *H2-hand*, the identity of the hand performing the second hand-move; *H2-target*, identity of the final target reached in the second hand-move. All the classification problems are two-levels problems, with classes *right* vs *left*. Results from the LDA run on the different combinations of classification problems and training sets provided estimates for the accuracy with which it is possible to predict the future moves based on the previous whole-body climbing kinematics, or subsets of it. LDA accuracies were therefore taken as proxy measures of coarticulation and were referred to as *coarticulation indexes*.

All classification problems were initially tested taking as predictor the whole-body kinematics across *Epoch1* (from start to *Hand1* onset), and independently across *Epoch1*+*Epoch2* (from start to *Hand2* onset). Note that for the first case, *H1-hand* is a depth-1 problem (predicting the next-to-come move) while *H2-hand* and *H2-target* are depth-2 problems (predicting the second-next-to-come move). For the second case, *H2-hand* and *H2-target* are depth-1 problems, while *H1-hand* is taken as method validation proof. In this case, in fact, the kinematics taken as predictor contains explicit information about *Hand1* (whose move unfolds during *Epoch2*) and therefore we expect virtually perfect classification performances.

In a second step of the analysis the LDA was run on smaller temporal chunks of the whole-body kinematics, with the aim of inspecting the temporal emergence of the coarticulation. In the last step of the analysis, the whole-body kinematics in each temporal chucks were further subdivided into single joints to further inspect which body parts coarticulate the most in the different stages of the climbing execution.

Running LDA on the whole-body kinematics (or subsets of it) spanning several seconds is however unfeasible due to the extremely high dimensionality of the input space, and the limited number of training trials. For this reason, it was necessary to derive compact descriptions of the whole-body kinematics during the various temporal intervals of interest.

#### Dimensionality reduction of whole-body climbing kinematics

To derive low-dimensional descriptions of the climbing kinematics we adopted spatiotemporal principal component decomposition (stPCA), a dimensionality reduction technique previously developed for the study of throwing behaviour (48,49). These previous works showed how stPCA allows to efficiently derive compact accurate descriptions of whole-body throwing actions: e.g., 10 (23) stPCs vectors are sufficient to account for 90% (97%) of the variance in a sample of over 2000 throws from 20 participants. The method could be applied to any multi-joints action unfolding over an extended time-interval. A crucial aspect of the method is that the kinematics of individual trials (action repetitions) should be time-normalized and resampled on a common temporal vector. When considering the case of climbing, it is important to take into consideration its sequential nature: it is indeed critical to temporally rescale trials so to align all the key-events that delimit the task segmentation. All trials from each individual participant were temporarily aligned so that the *onset* and the *reaching* event of the four end-effector moves included in the task (F1, F2, H1, H2) corresponds to the same temporal frame in the resampled timeline, as shown in Figure 2 (note that the *Climb Start* event is set to be the onset of the first foot-move, i.e. *F1-Onset*). More specifically, the whole climbing tasks was resampled on 180 frames, with each *move* phase (from *onset* to *reaching* of a given limb’s move) remapped onto 30 frames and each *adjustment* phase (from the *reaching* of a limb move to the *onset* of the subsequent limb’s move) mapped onto 20 frames. This choice of the resampled timeline roughly reflects the ratio between the average durations of the different phases of the climbing task between and across participants, while proving a sufficient granularity for inspecting the dynamics of the different stages of the sequenced route’s execution (see Figure 3B). It should be noticed that, despite the resampling procedure removes information about the average duration of each move and adjustment phase, the overall velocity profile of the joints kinematics is preserved, hence the information loss is minimal.

**Figure 3.**
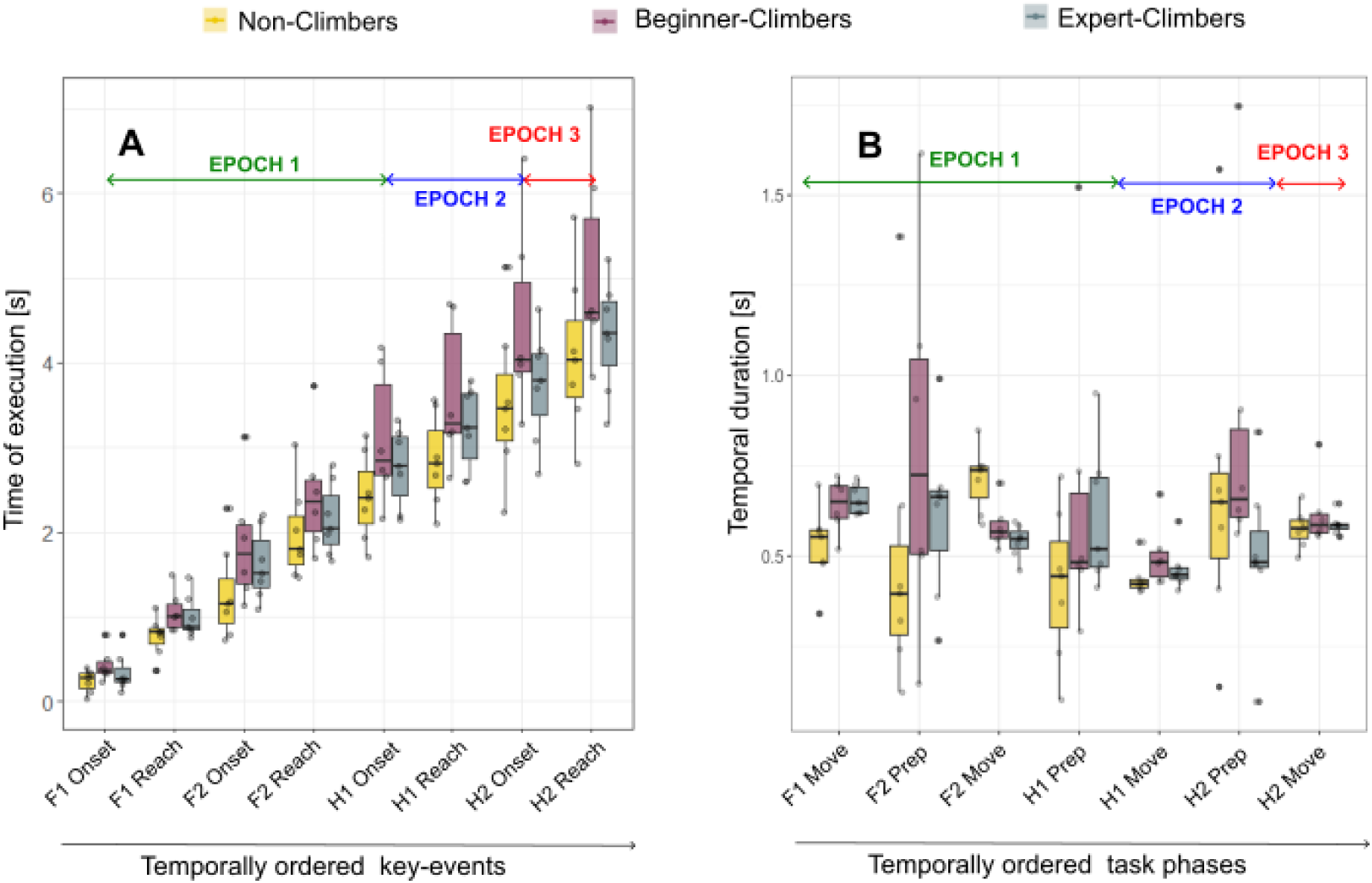
Temporal profiles of task execution across participants. (A) The distributions of the individual average times at which the key-events of the climbing occur are shown for participants belonging to the three expertise levels. (B) Distribution of the individual average duration of the different phases of the climbing task, shown for participants belonging to the three expertise levels. Points overlaid onto the boxplots represent the individual averages across trials repetitions. The boxplots show the distributions of the individual averages in each group.

In the different steps of our analysis, we were interested in assessing (i) whether climbers do coarticulate or not, (ii) how coarticulation emerges in time, and (iii) which body parts are the most involved in coarticulation. For this, we trained linear discriminant classifiers with different spatiotemporal subsets of the 3D positional data from the 25 recorded markers across the entire climbing task execution, deriving corresponding stPCs compact descriptions of the kinematics to be including in each training set. stPCA was therefore applied to obtain compact descriptions of (i) the whole-body encompassing *Epoch1*, and *Epoch1* plus *Epoch2*, (ii) the whole-body kinematics spanning the 18 temporal intervals in which the entire climbing execution unfolded (each remapped onto 10 timeframes), and (iii) the kinematics of each of the 25 tracked joints trajectories in each time bin.

The compact descriptions obtained were then used to train different classifiers and compute the individual *coarticulation index* for the whole-body kinematics across different time intervals. In addition, *coarticulation maps* providing the *coarticulation index* associated to each joint and temporal bin were derived for each participant. To quantify the extent of whole-body coordination, from the *coarticulation maps* we extracted the number of joints that in each temporal interval has a *coarticulation index* higher than 0.8, denoting it *N*_*Coart*−*Joints*_.

### Statistical analysis

In the statistical analysis we tested the effect of the climber’s expertise on the timings of the climbing execution and on the degree of coarticulation derived for the three classification problems tested in the LDA. The participant’s expertise was treated as a categorical factor, named *Expertise*, with three levels (*non-climbers, beginner-climbers, expert-climbers*). We further introduced other two main factors: the identity of key-events, named *i*_*KE*_, treated as an ordinal factor with values temporally ordered from 1 to 8, that was used for the timings analysis; the identity of the classification problem, named *P*_*i*_, treated as a categorical factor with levels *H1-hand*, *H2-hand*, and *H2-target*.

#### Timings of the climbing execution

Participants were free to execute the task at their own pace. Differences in the timing execution of the climbing task and of its different phases were tested. We extracted the times at which the 8 key-events occurred in each trial and took those values as the dependent variable in the Linear Mixed Model (LMM) described in Eq. 1. The model tests for the hypothesis that time at which each key-event occurs (*t*_*i*_) depends on *Expertise*, on the event itself (*i*_*KE*_), and on the subject identity (*ID*) – the latter treated as a random effect over trial repetitions.

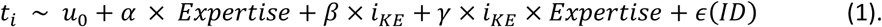

The duration of the 7 phases in which the climbing route’s execution was segmented were derived for each trial, and differences driven by *Expertise* were assessed with the LLM in Eq.2 applied to each phase independently.

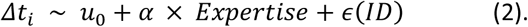

#### LDA classification and coarticulation index

LDA results are reported in terms of the classification accuracy taken as a proxy for the degree of coarticulation and used here to define the *coarticulation index* (*CI*). For each individual participant we computed the mean classification accuracy and the corresponding confidence interval obtained from a k-fold method with 20 folds – corresponding to 20 LDA repetitions in which the trials are divided into a training set of about 76 trials and a test set of about 4 trials (with little variations when among the 80 recorded trials some were excluded from the analysis). These results were used for generating the plots in Figures 6 and 7 for classification runs on the whole-body kinematics throughout *Epoch1* and *Epoch1+Epoch2*, and across the 18 time-bins encompassing the entire execution of the climbing task respectively. Statistical differences in *CI* across expertise levels and classification problems (*P*_*i*_) were tested with linear mixed models in all cases. Models were run on the extended dataset including all the 20 estimates of the individual coarticulation index resulting from the k-fold approach, so to implicitly account for the intrinsic accuracy of the individual *CI* estimates.

LDA results based on the whole-body kinematics throughout *Epoch1* and *Epoch1+Epoch2* were tested with Eq. 3, which assumes that the *Coarticulation Index* depends on *Expertise*, on the classification problem *P*_*i*_ (*H1-hand*, *H2-hand*, *H2-target*) and on their interaction, and including individual variability as a random factor ɛ.

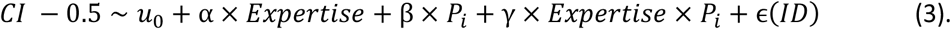

In Eq. 3 *CI* was modelled with an offset of 0.5 with the aim of testing the hypothesis that its values are above chance level, which corresponds to having a positive intercept in Eq. 3. In addition, one-tailed t-test were run on the distributions of *coarticulation indexes* obtained for specific combinations of *Expertise* and classification problem.

LDA results from the whole-body kinematics in the 18 time-bins in which the climbing task was segmented were tested independently for each classification problem with the following model:

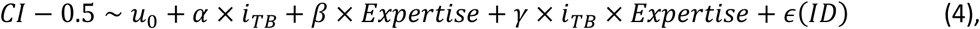

where *i_TB_* is the time-interval identifier treated as a categorical factor, and the other variables have the same meaning as in Eq. 3.

The number of joints that in each time-interval has a coarticulation index higher that 0.8, *N*_*Coart*−*Joints*_, was compared across participants with the model in Eq. 5, testing its dependence on *Expertise* and time:

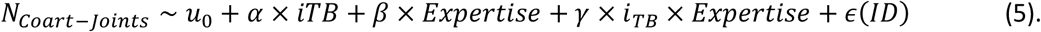

As for the other statical LMM tests the analysis was run on the sets of *N*_*Coart*−*Joints*_values obtained for each classification problem from the output of 20 LDA runs.

## RESULTS

Before testing the hypotheses under scrutiny, we inspected interindividual differences in the preferred stepping pattern, as well as in the timings of the execution of the whole task and of its different phases. In addition, we report on the efficiency of the stPCA in proving compact and accurate descriptions of the whole-body kinematics across extended time-intervals (i.e., *Epoch1* and *Epoch1+Expoch2*). The last part of the section focuses on testing the hypothesised *experience*-driven differences in the coarticulation indexes derived from the LDA.

### Stepping pattern and timings

The climbing task always started with a first phase in which both feet reached the two dedicated holds while grasping the initial hold, G_0_, with both hands. This passage is necessary for the reaching of G_1_ and G_2_ in the following part of the task. Participants were not instructed to comply with a fixed stepping sequence. Despite being all right-handed, they exhibited a clear trend for an individual preferred stepping sequence, with most participants consistently stepping first with the left foot: 14 out of 21 participants stepped with the left foot for more of the 93% of the trials; 3 participants showed an alternating behavior, with the first foot moved up swapping between right and left; only 3 participants consistently stepped first with the right foot (for more than 98% of the trials).

The individual average time at which the key-events occur, *t_KE,_*are shown in Figure 3A, with participants grouped according to the three levels of *Expertise*. A clear trend is visible: *beginner-climbers* tend to take more time, with respect to both *non-climbers* and *expert-climbers*, to execute all stages of the climbing execution. Statistical significance of such effect was tested using the Linear Mixed Model (LMM) described in Eq. 1. The LLM model assumes that the time at which key-events are executed increases with the *i_KE_*, with *i_KE_*defined as an ordered numerical factor that identifies the key-events in their temporal sequence. The model further assumes that this dependence is modulated by the level of expertise, i.e. *Expertise* treated as a categorical factor. LLM results confirm a strong dependence of *t_KE_* on *i_KE_* (*p* < 10^−10^), and a significant effect of *Expertise*, where the fitted slope is significantly higher for *beginner-climbers* than for *non-climbers* (p = 0.021), and tends to be higher with respect to *expert-climbers* (*p* = 0.13). The model output details are reported in Table S1. Note that the lack of significant interactions between *i_KE_*and *Expertise* implies that the effect of *Expertise*, reported as main effect, is significant at all *i_KE_*.

To inspect how these differences cumulate across the various phases of the climbing task we looked at the distributions of their durations in the three *Expertise* groups (Figure 3B). Running the LMM model in Eq.2 for each phase independently, we found a significant effect of *Expertise* only in the extent of the first foot-move and second foot-move duration that were respectively longer and shorter in *non-climbers* than in *beginner-climbers*. There are however a few trends for the phase durations being longer in *beginner-climbers* (see Figure 3B) that, by cumulating across subsequent phases, can explain the main effects of *Expertise* reported above.

### Low-dimensional descriptions of the whole-body kinematics in a climbing task

Spatiotemporal Principal Component Analysis (stPCA) (48,49) was applied independently for each participant, for the whole-body kinematics recorded over extended temporal durations across the climbing task (*Epoch1* and *Epoch1*+*Epoch2*), and for different subsamples of it in time and body space.

When applied to the whole-body kinematics over *Epoch1* and *Epoch1+Epoch2*, stPCA could achieve highly accurate descriptions of the recorded whole-body kinematics as combinations of a small number of components. For example, the 3D positional data of the 25 markers sampled onto 100 (for *Epoch1*) or 150 (for *Epoch1*+*Epoch2*) time samples could be reconstructed as a combination of 15 components (stPCs) vectors while accounting for above 93% of the total variation in each individual dataset; values above 99% of the total variation could be achieved using up to 40 stPC vectors for most participants (but 2 over 20). Considering that the initial input space has a dimension above 7500, these numbers provide clear evidence of the efficacy of stPCA in achieving highly accurate descriptions with a drop of more than two orders of magnitude in dimensionality.

Figure 4 shows a representative example of how well the recorded kinematics (shown in blue) can be approximated by the compact sPCA descriptions (shown in red), and how the reconstruction accuracy progressively increases when incorporating higher order stPC vectors (from 10, to 20, to 40). When applied to temporal and spatiotemporal (in body space) subsamples of the whole-body kinematics, high reconstruction accuracies could be achieved with a smaller number of stPC vectors. For the reconstruction of the whole-body kinematics each across the 18 time-intervals we used 25 stPCs, while for the reconstruction of single-joint kinematics in each time-interval we used 5 stPCs.

**Figure 4.**
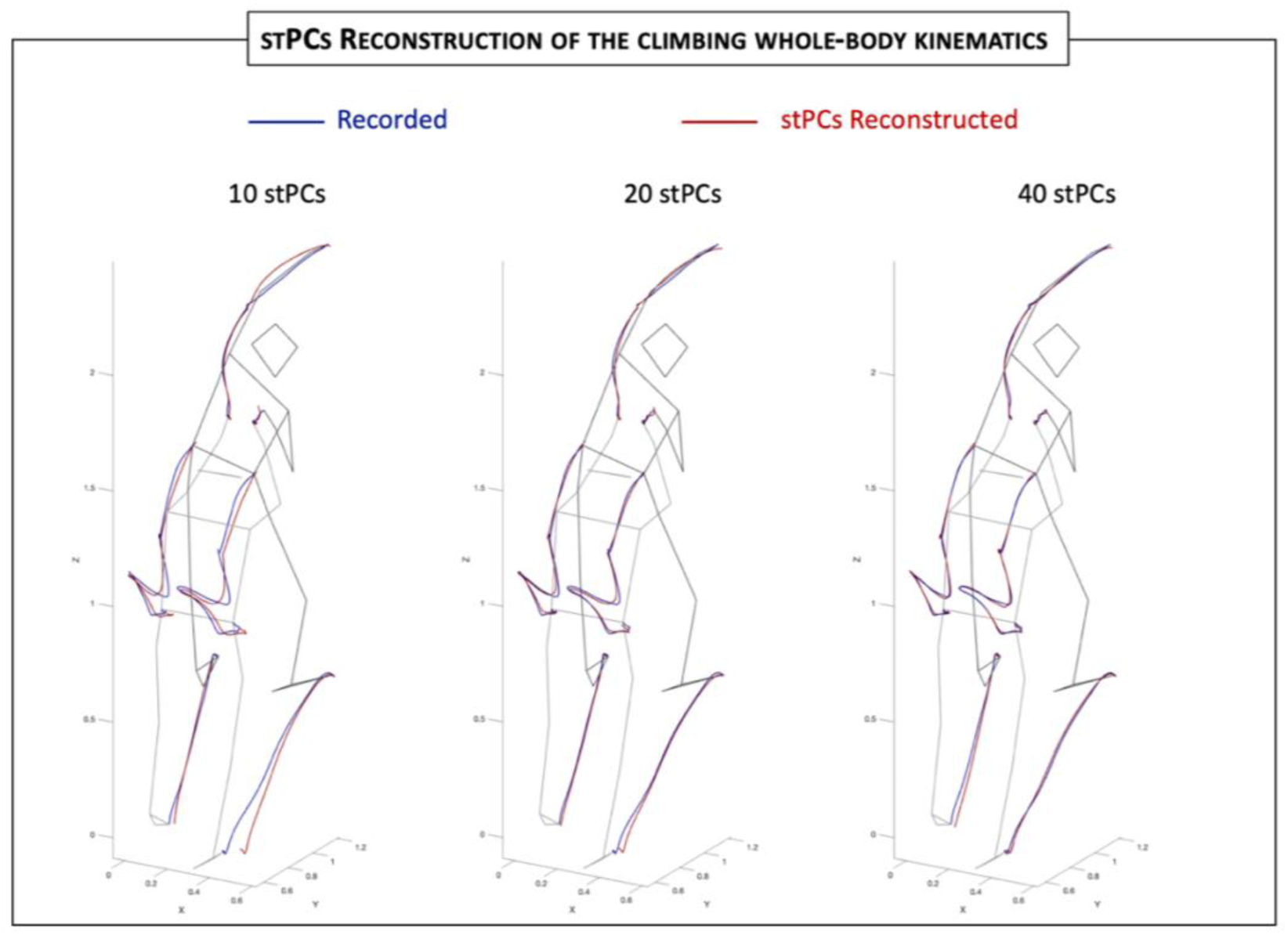
Real and stPCs-reconstructed whole-body kinematics. The red trajectories show the kinematics of the end effectors (hands and feet) and of the two hip joints reconstructed with 10 (left), 20 (centre) and 40 (right) stPC vectors, compared with the real recorded kinematics shown in blue, for the case of a single trial from subject S20. The two stick diagrams in each panel shows the body posture at the beginning of the trial and the end of Epoch 2.

### Coarticulation in climbing kinematics

The choice of climbing as an intrinsically sequential motor task interlaced with planning was tuned to the aim of assessing coarticulation in naturalistic whole-body motor behaviours. To this aim the climbing tasks was segmented and the *coarticulation index* (*CI*) was derived using LDA based on the whole-body kinematics, or single-joint kinematics, across different time-intervals encompassing various phases of the task sequence.

### Whole-body kinematics provide signatures of coarticulation

As a first step we explored coarticulation at the individual level, estimating the extent to which the whole-body kinematics during the task execution (across *Epoch1* and *Epoch1+Epoch2*) can predict the action(s) that will be performed next (e.g., which hand will be moved next, and to which target hold). To quantify the degree of coarticulation we run LDA for the three classification problems of interest: *H1-hand*, *H2-hand*, and *H2-target*. LDA classifiers were trained with the whole-body kinematics descriptions obtained from the linear combination of the first 35 stPC vectors. This choice was based on a preliminary exploratory analysis in which we compared LDA results obtained with stPC whole-body kinematics reconstructions with dimensionalities ranging from 15 to 50. Results with 35 stPCs are shown in Figure 5. The *coarticulation indexes* (*CI*s) of individual participants are shown with the corresponding confidence intervals as the circular markers and shaded error bars in Figure 5, with the distributions of the individual average values in the three groups of *Expertise* shown by the overlayed boxplots. Results from LDA classification trained with the whole-body kinematics throughout *Epoch1* and *Epoch1+Epoch2* are shown in Figure 5A and 5B respectively.

**Figure 5.**
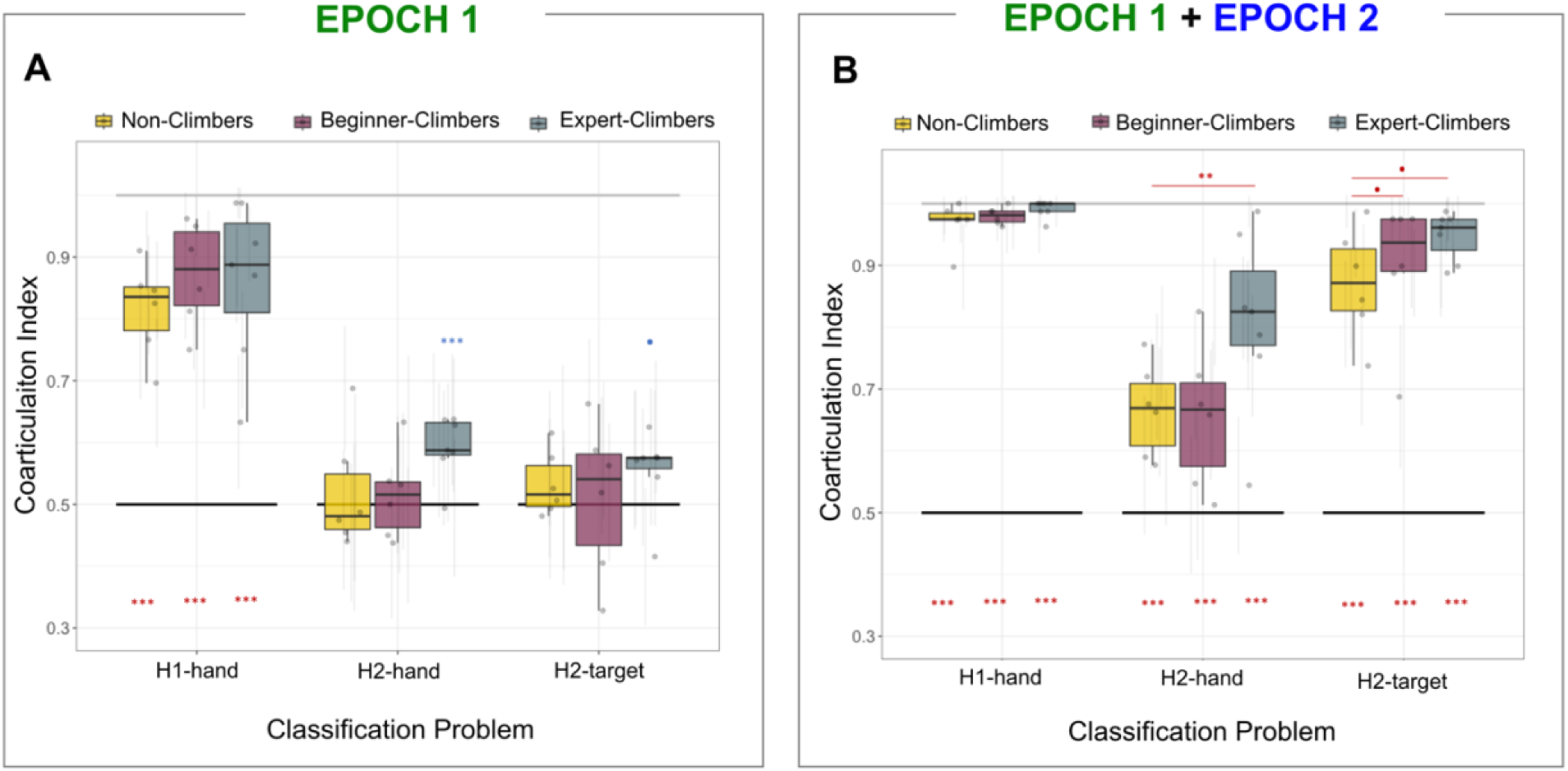
Degree of coarticulation during Epoch1 and Epoch 2. The distribution of the *coarticulation indexes* across participants with different levels of climbing expertise is shown for different classification problems, classifying for: *H1-hand*, i.e. the hand (left-vs-right) performing the first hand-move; *H2-hand*, i.e. the hand performing the second hand-move; and *H2-target*, i.e. the target (left-vs-right) of the second hand-move is directed. The boxplots show the distributions of the individual averaged *coarticulation indexes* averaged across repetitions, defined as the classification accuracy obtained with a k-Folds validation approach (20 folds); the light shaded vertical lines showing the corresponding confidence interval. The horizontal lines at 0.5 (*black*) and 1 (*grey*) indicate the reference for chance level and perfect classification performance.

Differences in *CI* values across expertise level (*Expertise*), classification problems (*P*_*i*_) and their interaction were analysed with the linear mixed model in Eq. 3, run on the extended dataset including the classification accuracies obtained in all the 20 folds LDA repetitions. The detailed LLM outputs are reported in Table S1 and Table S2. In the following we discuss the most relevant results.

Taking the whole-body kinematics throughout *Epoch1* as predictor we obtained the following outcomes. First, we found a significant degree of coarticulation driven by the first moving-hand: *CI* values for the *H1-hand* problem are significantly above 0.5 (i.e., chance level for correct classification) for *expert-climbers* (*p* < 10^−10^, *mean* ± *std* = 0.86 ± 0.03) and not significantly different from the other two *Expertise* groups, which are indeed above chance level (*beginner-climbers*: *p* < 10^−10^, *mean* ± *std* = 0.87 ± 0.03; *non* − *climbers*: *p* < 10^−10^, *mean* ± *std* = 0.81 ± 0.03). For all groups the accuracy in the *H1-hand* problem is found to be significantly higher than for the other two classification problems *H2-hand* and *H2-target*, which both corresponds to coarticulation at “depth-2”, meaning that – unsurprisingly – the degree of coarticulation is higher for the immediate following next-to-come move than for the more distant second next-to-come move. No significant interactions between *P*_*i*_ and *Expertise* is found, implying that the degree of coarticulation is not significantly modulated by expertise also in *H2-hand* and *H2-target*. Still in Figure 5A it could be appreciated how the degree of coarticulation in *expert-climbers* tend to be above chance levels in both *H2-hand* and *H2-target*, which is not the case for the other two expertise groups. To better inspect this, we run one-tail t-tests for each *Expertise* group in the *H2-hand* and *H2-target* problems. After Bonferroni correction, we found that the distribution of *CI* values for *expert-climbers* is significantly above chance level only for *H2-hand* (*p* = 2 × 10^−4^), although with a low accuracy (*mean* ± *std* = 0.59 ± 0.03), while for *H2-target* a trend for being above chance level was found for both *expert-climbers* (*p* ≈ 0.11) and *non-climbers* (*p* ≈ 0.054). No evidence for coarticulation was found for the other cases. In summary, these results show that there is a significant degree of coarticulation at depth-1, i.e. with respect to the hand that will perform the next-to-come movement for all participants, while only some *expert-climbers* exhibit some degree of coarticulation at depth-2 (i.e., with respect to the second next-to-come hand-move), mainly driven from which of the two hand (right or left) will perform the move. Weak evidence for coarticulation with respect to the target of the second hand-move was further found in both *non-climbers* and *expert-climbers*.

Similarly to what discussed above, we assessed coarticulation in the whole-body kinematics that encompasses *Epoch1* and *Epoch2*, i.e. from climb start to the onset of the second hand-move. The results are show in Figure 5B with the same format adopted for Figure 5A. The LLM run on the LDA individual accuracies from the 20k-fold LDA approach highlighted several statistically significant results. First, *coarticulation index* is significantly above chance for all problems independently of the climber expertise: the *H2-hand* problem, taken as the reference value in the mixed model, is significantly above chance level for the *non-climbers* group (*p* ≈ 2.6 × 10^−7^; *mean* ± *std* = 0.67 ± 0.02); no significant difference is found between *non-climbers* and *beginner-climbers* (*mean* ± *std* = 0.63 ± 0.04), while *expert-climbers* are characterized by significantly higher *CI* values (*mean* ± *std* = 0.80 ± 0.03) with respect to *non-climbers* (*p* ≈ 1.2 × 10^−3^). *CI* values in *non-climbers* are significantly higher for the *H2-target* than for the *H2-hand* (*p* < 10^−10^; *mean* ± *std* = 0.87 ± 0.02), and the *H1-hand* classification problems (*p* < 10^−10^; *mean* ± *std* = 0.98 ± 0.02). Note that the latter case is to be regarded as a validation of the adopted methodology, because the predictors contain explicit information of which hand performs the first hand-move (during *Epoch2*). The model further highlights that for *H2-target* there is a trend for the classification accuracy for both *beginner-climbers* (*p* ≈ 0.10; *mean* ± *std* = 0.90 ± 0.03) and *expert-climbers* (*p* ≈ 0.13; *mean* ± *std* = 0.95 ± 0.02) to be higher than for *non-climbers*. In summary, these results show that i) all participants coarticulate their whole-body kinematics throughout *Epoch 2* with respect both to which hand (right or left) will move next and where to, ii) the coarticulation driven by the target location is stronger than the one driven by which of the two hands will perform the move, iii) climbers display a higher degree of coarticulation with respect to participants with no climbing experience.

The results shown so far provide evidence for coarticulation to be present to some extent in the whole-body kinematics for all participants. In addition, results show that the degree of coarticulation is modulated by the level of climbing expertise. The previous analyses however do not provide information about how such coarticulation is structured in time and which are the motor strategies are employed. In the following we analyse these aspects.

### Temporal unfolding of coarticulation

To test the temporal unfolding of the degree of coarticulation, the whole-body kinematics recorded during the climbing tasks has been divided in 18 time-intervals. LDA was then run on the 25-dimensional stPCs descriptions of these subsets of whole-body kinematics. For each participant and each time-interval LDA was run independently.

Figure 7 shows the temporal unfolding of the *coarticulation indexes* extracted for the three classification problems (*H1-hand*, *H2-hand* and *H2-target*), highlighting individual trends (points connected by thin lines), as well as summary trends for the three *Expertise* groups. Differences across groups and time-intervals were tested with the linear mixed model described by Eq. 4 and run independently on the three classification problems. The detailed outputs of all three LLMs are given in Tables TS4 –TS6. In the following we discuss the most relevant results.

For the *H1-hand* problem, coarticulation was on average above chance level in all time-intervals, independently on *Expertise.* The estimate for average coarticulation in the *non-climber* group for the first interval is significantly above chance (*p* = 6.8 × 10^−4^ and *mean* ± *std* = 0.67 ± 0.04), with no significant difference from other groups. As can be seen in Figure 7A, this results from averaging two main trends observed at the individual level: some individuals “coarticulate” since the beginning of the climbing task, positioning themselves differently according to the hand they will have to move first; other participants start instead from a neutral position, not informative of the planned next move. Importantly, these two main trends were not associated with *Expertise*. In the following time intervals, the *CI* starts to be significantly and consistently higher with respect to its initial value from the 8^th^ time-interval, which corresponds to the *adjustment phase* between the reaching of the second foot (*F2-*reach) and the onset of the first hand (*H1-*hand). No significant difference across expertise groups is found, beside for the *beginner-climbers* group characterized by lower *CI* values that the other two groups in the 8^th^ time-interval.

For the *H2-hand* classification problem, *CI* values for *non-climbers* start to be significantly and consistently above chance level (*p* < 10^−8^ and *mean* ± *std* = 0.74 ± 0.03) at the 14^th^ time-interval, corresponding to the latest stage of the “adjustment phase” preceding the second hand move, with no significant difference from other groups. Looking at interaction effects, it emerges that *beginner-climbers* display significantly larger *CI* with respect to *non-climbers* (*p* < 0.04) in the 16^th^ and 17^th^ time-intervals, namely during the second hand-move. *Expert-climbers* instead display significantly larger *CI* values with respect to *non-climber* starting from the 11^th^ time-interval forward (*p* < 0.04), which is already during the first-hand move.

Results from the *H2-target* problem reflect a higher degree of coarticulation with respect to *H2-hand.* In this case, *CI* values for the *non-climber* group start to be significantly above chance level earlier, at the 11^th^ time-interval (*p* < 10^−4^ and *mean* ± *std* = 0.62 ± 0.03), beside the isolated case of the 1^st^ time-interval (*p* < 10^−4^ and *mean* ± *std* = 0.62 ± 0.03). At the 11^th^ time-interval, in the initial stage of the first-hand move, *CI* gets significantly above chance level for the *non-climber* group (*p* = 0.016 and *mean* ± *std* = 0.60 ± 0.03) and increases progressively thereafter. Since the 11^th^ interval *CI* values are significantly higher in *expert-climbers* that in *non-climbers*. No significant differences between *non-climber*s and *beginner-climbers* are found, but for *CI* values being significantly lower in *beginner-climbers* in the 9^th^ and 10^th^ intervals, i.e. during the adjustment phase preceding the onset of the second hand-move.

In summary, we found that all climbers in our sample display some degree of coarticulation, although *expert-climbers* tend to engage in preparatory behaviour at earlier stages and, differently from the other groups, show evidence for coarticulation at depth-2.

**Figure 6.**
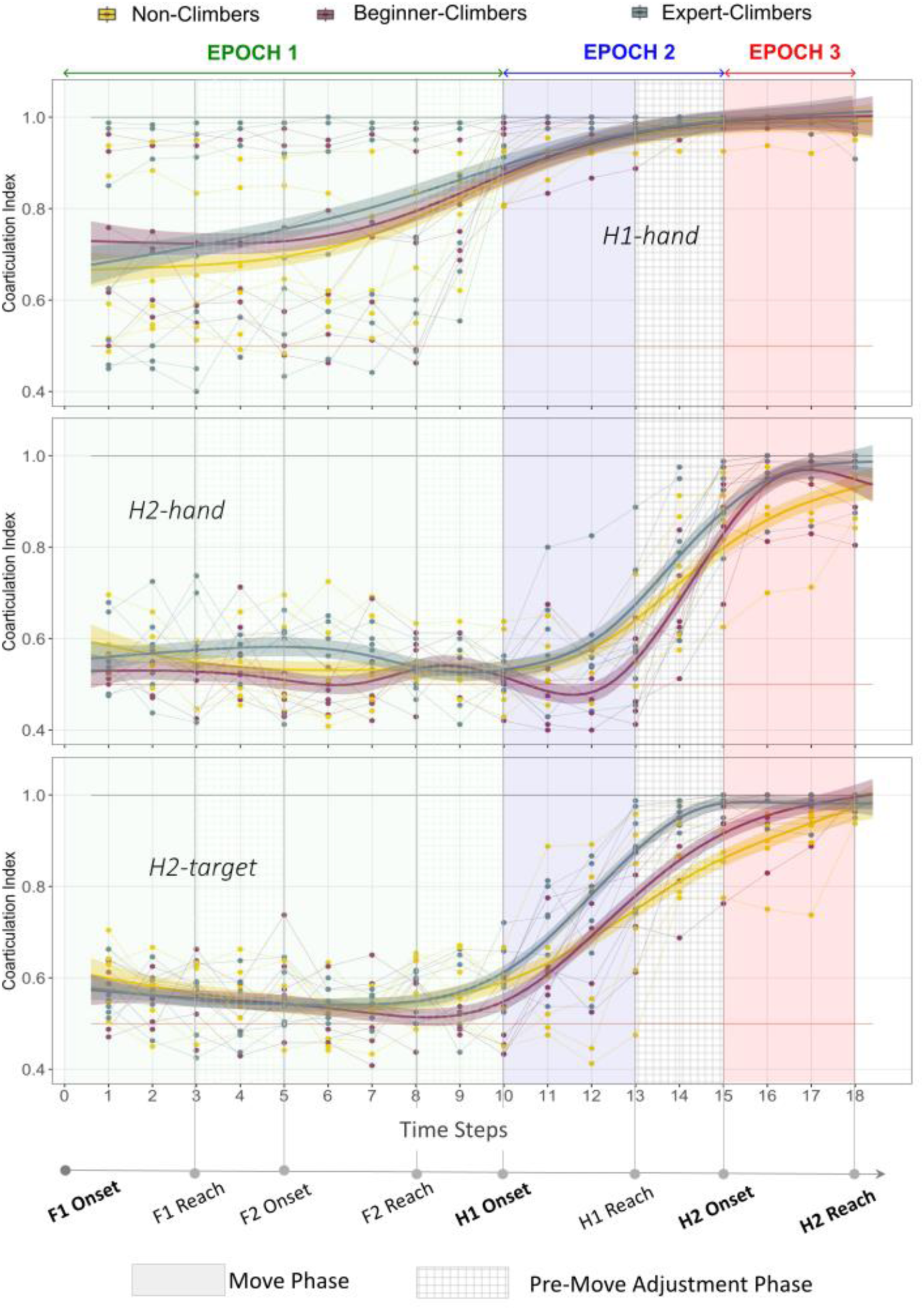
Temporal unfolding coarticulation throughout the climbing task. The figure shows the individual average *coarticulation indexes* at 18 time-intervals spanning the whole climbing task, as point data coloured by *Expertise*; thin lines connect the points associated with individual participants. The continuous lines and the overlaid shaded areas show smoothed conditional means and associated confidence intervals, as computed by the *geom_smooth* function in R, applied to the three *Expertise* groups, and are shown as an aid to highlights patterns in the data.

### Coarticulation in body space

As a last step in our analysis, we aim at characterizing how coarticulation is operated in terms of motor strategies, by identifying the body parts engaged in the coarticulation at the different phases of the climbing tasks. To this aim, we applied stPCA + LDA to the chunks of single-joint trajectories obtained with the same temporal segmentation used above. The output of this analysis returns individual coarticulation matrixes, providing the *Coarticulation Index* as a function of the tracked body joints and of the temporal unfolding of the climbing task execution.

Figure 7 shows the results from the three classification problems under analysis for a representative *beginner-climbers* participant. The participant belongs to the group that coarticulate from the very beginning: this can be appreciated from the left panel, displaying results from the *H1-hand* problem, where the two red horizontal bands corresponds to the two arms and indicate how the subject adapts her initial posture modifying the configuration of the arms according to the hand that will perform the first hand-move. Comparing the maps in the central and right panels, it is clearly visible that the degree of coarticulation is higher for *H2-target* than for *H2-hand*: the subject exhibits a depth-2 coarticulation driven by the final target location at the very beginning of the task, which vanishes before the second foot-move and reappears during the adjustments preceding the second hand-move. Crucially, the spatiotemporal patterns of coarticulation for the different participants exhibit a high degree of idiosyncrasy which does not exhibit any obvious relation with expertise. For space issues coarticulation maps from all participants are provided in the Supplementary Material (Figure S1-S3).

Differences across participants driven by *Expertise* were tested counting the number of joints, *N*_*Coart*−*Joints*_, that in each time-interval has a *Coarticulation Index* above 0.8. Results are shown in Figure 8 for the three classification problems of interest. *N*_*Coart*−*Joints*_ was computed as a proxy for the degree of whole-body coordination involved in individual coarticulation strategies, and its dependence on *Expertise* across the different phases of the climbing task was tested running the LMM in Eq. 5 for each classification problems. The complete outputs of the models are given in Tables TS7-TS9. Below we discuss the most relevant results.

For the *H1-hand* problem *N*_*Coart*−*Joints*_ displays the expected overall increase with time. Significant differences driven by *Expertise* emerges progressively: *N*_*Coart*−*Joints*_ results to be significantly higher in *beginner-climbers* than *non-climbers* starting from the adaptation phase preceding the second foot-move (*p* = 0.0054 for the 5^th^ time-interval, and *p* < 10^−3^ for intervals ranging from 6^th^ to 14^th^). At the same time *expert-climbers* are characterized by significantly larger *N*_*Coart*−*Joints*_ than *beginner-climbers* during the adaptation phase preceding the first hand-move (*p* = 0.045 and *p* < 10^−6^ for the 9^th^ and 10^th^ time-intervals respectively), and during its initial stage (*p* = 0.0093 for the 11^th^ time-interval).

Significant differences driven by *Expertise* are also found in the *H2-hand* and *H2-target* problems. In the first case, *N*_*Coart*−*Joints*_ results to be significantly higher in *beginner-climbers* than *non-climbers* starting from the adaptation phase preceding the second foot-move (*p* < 0.05 for the 4^th^ to 9^th^ time-intervals) all the way through the end of the task execution (*p* < 0.02 for all intervals but the 9^th^ and 15^th^ in no significant difference is found). *N*_*Coart*−*Joints*_ is found to be higher in *expert-climbers* than in *beginner-climbers*, starting from the onset of the first hand-move (*p* = 0.03 for the 11^th^ time-interval) all the way through the end of the task (*p* < 0.002). The *H2-target* problem shows a different pattern of results. As in the other cases, *beginner-climbers* are characterized by higher *N*_*Coart*−*Joints*_ values than *non-climbers*, starting from the adaptation phase preceding the onset of the second hand-move (*p* < 10^−4^). However, in this case *N*_*Coart*−*Joints*_ results to be significantly higher in *beginner-climbers* also with respect to *expert-climbers* during the adaptation phase preceding the second hand-move onset (*p* < 0.02) and its execution (*p* < 10^−10^).

In summary this analysis shows a very large heterogeneity in the individual motor strategies adopted during coarticulation, which is observed both across and within levels of climbing expertise. Still, despite this huge variability, data show a general trend for *Expertise* to modulate the number of body joints characterized by a high degree of coarticulation, *N*_*Coart*−*Joints*_. The latter was introduced as a proxy for quantifying the extent to which coarticulation involves the coordination of multiple joints. In this respect, results show how *climbers* tend to modulate their motor execution with a higher degree of whole-body coordination with respect to *non-climbers*, an effect more pronounced in *expert-climbers* than in *beginners*.

**Figure 7.**
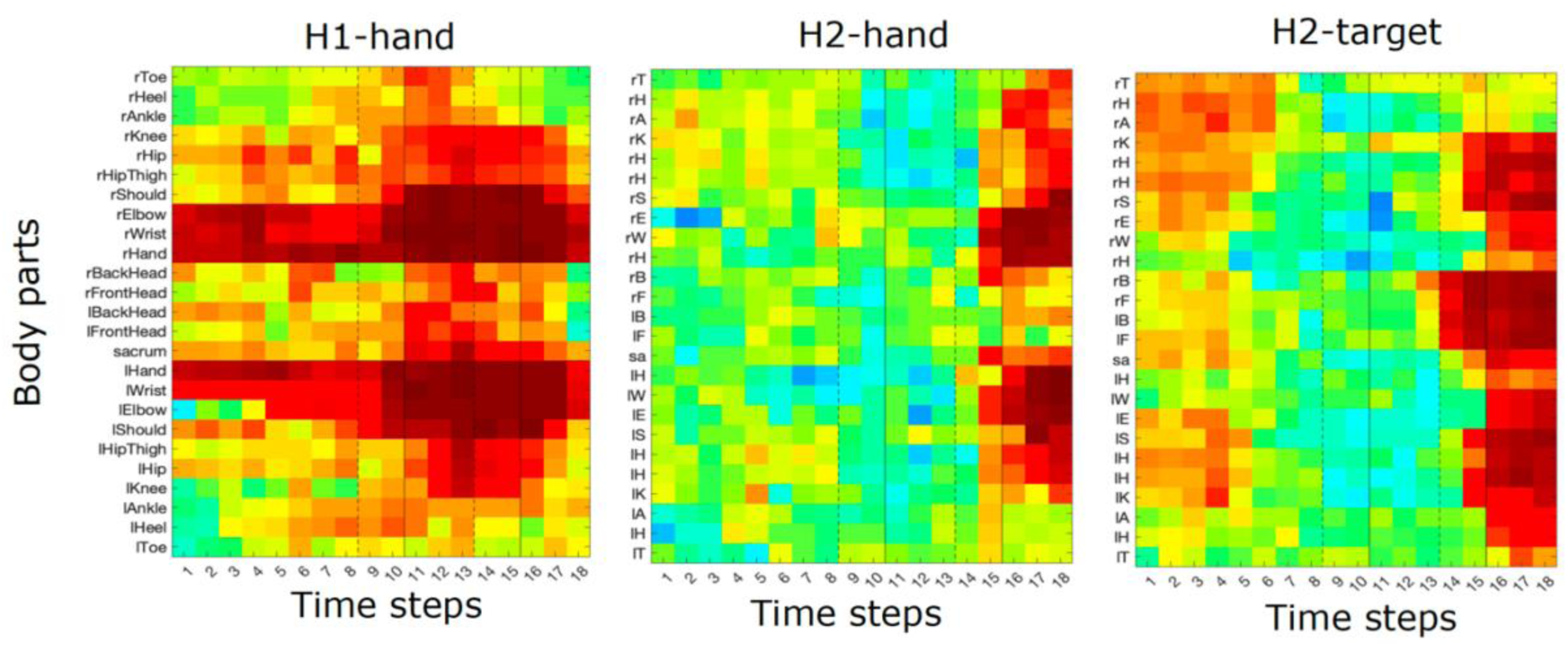
Individual spatiotemporal patterns of coarticulation indexes. The heatmaps depicts the coarticulation index as a function of the tracked body joints and of the temporal unfolding of the climbing task execution, for a representative *beginner-climber* participant and for the three classification problems predicting the hand (right or left) performing the first and the second hand-moves (*H1-hand* and *H2-hand*, left and central panels, respectively) and the target of the second hand-move (*H2-target*, right panel).

**Figure 8.**
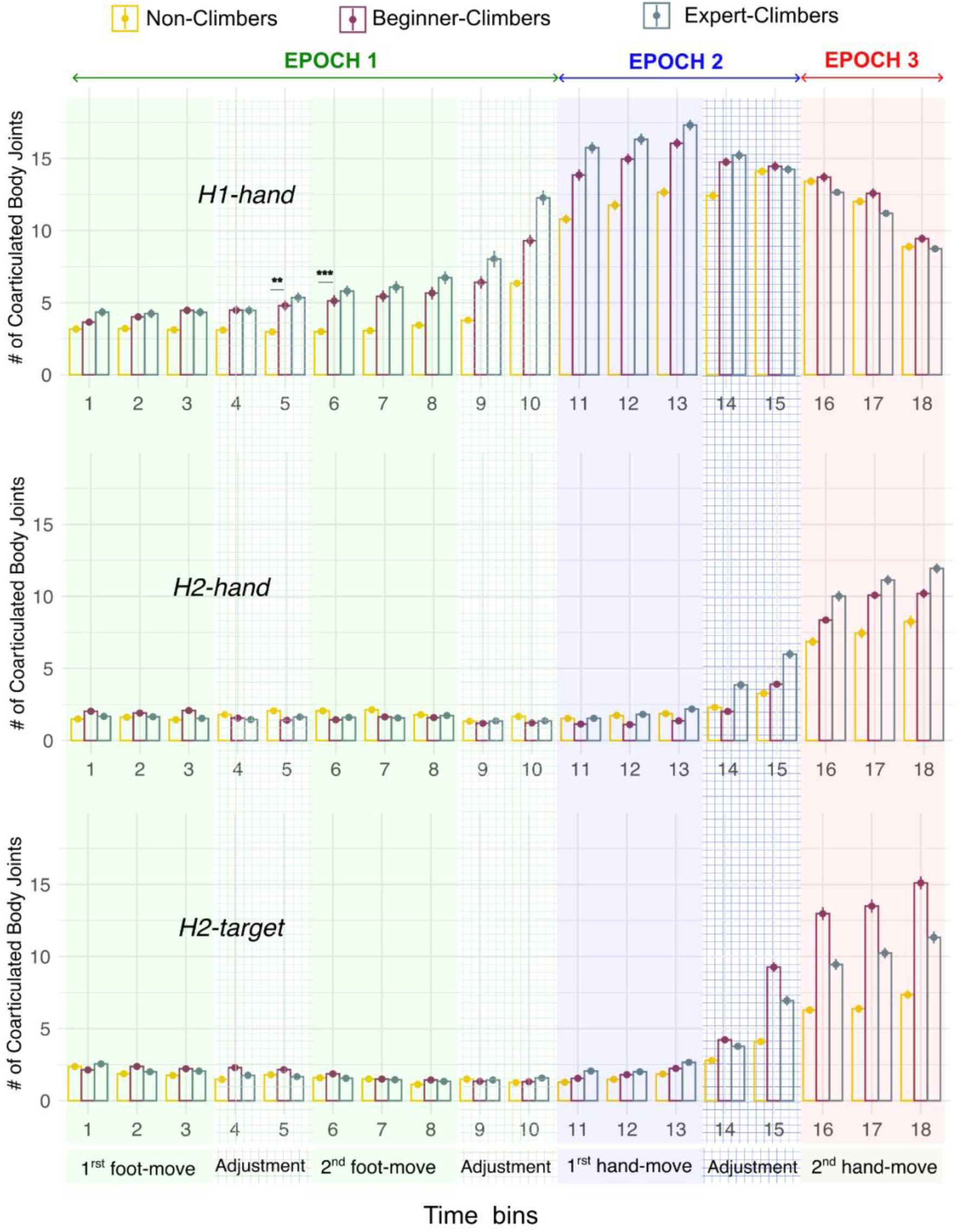
Number of highly coarticulating joints throughout task execution. The plots show the average number of joints with a *Coarticulation Index* higher than 0.8 (*N*_*Coart*−*Joints*_), for the three classification problems as indicated in each of the three panels. For each of the 18 temporal intervals, the individual *Coarticulation Index* was computed for all the 25 joints, to derive *N*_*Coart*−*Joints*_. The bar plots, shown for each time interval and for the three *Expertise* groups, represent the mean and standard error of *N*_*Coart*−*Joints*_values obtained running 20-fold LDA for each participant belonging to the same *Expertise* group.

## DISCUSSION

Sport climbing offers an ideal testbed to study the interplay between planning and skilled motor behavior (6). When facing a climbing problem, expert climbers need to first form a plan about the sequence of actions that could solve the assigned route, then they need to execute the route concatenating the planned sequence of whole-body movements. In this study, we build up on this specificity of sport climbing with the aim to provide a quantitative assessment of the impact of having a plan on naturalistic motor behavior. In our experiment the climbing sequence was instructed, so that the planning phase was limited to the memorization of the instructions given at each trial; in this way, we ensured that all participants had formed a clear plan before starting the execution, independently of their expertise. The aim of the study was to assess whether the formation of a climbing plan induces coarticulation in the motor execution of a set of climbing routes, and whether the degree of coarticulation depends on climbing expertise.

We report two main results. First, our analysis of the whole-body kinematics during the initial stage of the routes, from climb start to the onset of the first (or second) hand-move, shows that most participants engage in coarticulation to some extent, regardless of their expertise. Results show coarticulation to be significantly above chance for depth-1 problems in all three expertise groups. A new finding is that the the coarticulation driven by the location to be reached in the next-to-come move was found to be stronger than the one driven by the hand that has to perform the move. This might potentially reflect the fact that spatial aspects of the climbing problems, like the direction of the next hold to be grasped and the possible whole-body weight shift involved, are key determinants of whole-body movements coordination and coarticulation.

Second, we found that climbing expertise modulates coarticulation. Specifically, *expert-climbers*, but not *beginner-climbers or non-climbers*, modulate their climbing execution depending on the second-to-come move, i.e. display some degree of coarticulation also for depth-2 problems. Furthermore, our results show that *expert-climbers* exhibit a higher degree of coarticulation in those depth-1 problem for which coarticulation in non-experts is limited, as for the case of the coarticulation driven by the hand that will have to perform the second hand-move. Inspecting the temporal unfolding of coarticulation indexes provided additional insights on how expertise modulates coarticulation. *Expert-climbers* tend to engage in motor coarticulation at earlier stages with respect to climbers in the other groups. Specifically, *non-climbers* tend to prepare for the next move, mostly in the interval between its onset and the end of the previous move, which consists in posture adjustments with both feet and hands staying at place; *expert-climbers,* and to a less extent *beginner-climbers*, starts instead coarticulating during the execution of the previous hand-move. Finally, inspecting how coarticulation emerges in body space brought further interesting outcomes. A large heterogeneity in the individual motor strategies adopted during coarticulation was found both across and within levels of climbing expertise. Still, climbers (both experts and beginners) tend to engage in a higher degree of whole-body coordination with respect to *non-climbers*, as highlighted by the higher number of individual joints displaying coarticulation above chance-level; the effect was found to be more pronounced in *expert-climbers* than in *beginner-climbers*.

There results offer novel evidence for coarticulation in whole-body naturalistic motor behaviour, showing how the execution of a specific motor task, e.g. when having to move one hand from a hold to another while keeping the other hand and both feet fixed, is modulated according to what comes next in the plan. This adds to the rich repertoire of sequential motor behaviours in which coarticulation has been reported, ranging from speech production to finger tapping, extending it to encompass naturalistic whole-body movements and moving away from the constraints of laboratory settings (10).

This advance was achieved thanks to our methodological approach combining standard classification algorithms with stPCA, a dimensionality reduction technique specifically developed for dealing with complex whole-body kinematics (48,49). The use of stPCA was particularly crucial: while other methods, such as standard PCA (52) and the kinectome (53) have been proposed to cope with the high dimensionality of the whole-body kinematics avoiding the preselection of a limited number of movement features, these methods have been proved successfully mainly for cyclic motor tasks like walking, running or skying (54,55). Instead, stPCA allows to achieve compact descriptions of whole-body kinematics of open-ended actions spanning extended time-intervals. These low-dimensional descriptions allow for the comparison of motor performance across experimental conditions, within and between participants, with no need for collecting a huge amount of trial repetitions that would be otherwise necessary (56), but unfeasible in controlled laboratory settings.

Our findings add to a large literature on coarticulation, showing it for the first time at a whole-body level in a naturalistic task (sport climbing) and as a function of expertise. The effect of expertise in driving different degrees of motor coarticulation could have origin in different aspects of motor learning and motor control. In the context of motor learning, coarticulation studies have shown that practice and learning enhance coarticulation. The established view relates coarticulation to chunking, a process observed in tasks requiring a concatenation of multiple steps that consists in forming an operational representation of the task execution as a sequence of contiguous movement units (57,58). This segmented representation emerges as a cognitive strategy to facilitate the recalling processing required in complex compound tasks, but it correspond to a sub-optimal efficiency of their motor execution, which results in a series of stops and goes. With practice however, contiguous chunks tend to be merged resulting in smoother and faster motor execution and in fewer chunks displaying coarticulation (38,59): the execution of each chunk is progressively modulated to prepare and thus facilitate the execution of the following one(s), resulting in some cases in the fusion of two or more chunks into a single unit with no intermediate stops.

In line with chunking theories, the higher degree of coarticulation we found in *expert-climbers* may result from an enhanced capability in turning the abstract plan initially assigned as a sequence of discrete end-effectors’ moves, into a motor plan of how to execute such moves at the level of whole-body coordinated movements. This may imply an initial mapping of the discrete goal for the end-effector onto a whole-body target configuration to which the whole-body shall converge. In computational motor control, this mapping is believed to be supported by internal forward models (60,61) able to simulation whole-body movements to accomplish the plan. This control mechanism has also been suggested to underly the motor-cognitive planning processes in climbing (18). In addition, expertise is expected to enhance one’s ability to optimize the whole execution by modulating each step so to facilitate the rest of the task execution. This could result from the ability to select among the possible motor plans to the same intermediate discrete goal (e.g. right hand on hold G_1_), the one that minimizes the effort later required for completing the whole task.

Taking the perspective of motor control theories, the higher degree of coarticulation found expert-climbers might reflect a more refined optimization process that minimizes the overall cost of the climbing sequences execution (37,62). This is in line with complementary evidence from the literature showing how sport expertise correlates with more efficient visual behavior: experts exhibit longer fixations to task-relevant features of the task space and fewer fixations to task-irrelevant features with respect to novices and beginners (63). This overall behavior has been explained in the context of action selection processing (64,65) as a sort of “shielding mechanism”. Shielding occurs through inhibiting the processing of non-optimal task solutions and gets more efficient with practice and learning, which leads to the eventual selection and execution of optimized solution (66,67). Although these observations do not provide direct evidence for the recruitment of internal forward models guiding motor tuning, spending more time processing feasible solutions may result from the need to estimate possible alternative motor strategies together with their corresponding overall efficiency, a process that would indeed need the recruitment of forward models. Current data are nevertheless not sufficient to substantiate this speculation and future studies are needed to investigate further this possibility.

The large inter-individual variability found in how coarticulation emerges across different body parts during the task execution reflects the large inter-individual variability that characterizes the execution of complex multi-joints motor tasks, which are intrinsically redundant and can be accomplished with different motor strategies (7,49,68–70). Beside this expected variability, we found a systematic trend for experts engaging in higher degrees of whole-body coordination. This result fits well within the established evidence of the progressive emergence of patterns of highly coordinated whole-body movements during motor development and motor learning (71–75).

While providing one of the first evidence for coarticulation in naturalistic whole-body motor behavior, the current study has few limitations that should be addressed in follow-up works. First, the climbing routes assigned to our participants were short and their execution did not require advanced motor skills. While this choice ensured that non-climbers would be able to execute the required high number of routes repetitions, it limited the extent to which coarticulation can emerge and be observed. This implies that our procedure might have led to an underestimation of the degree of coarticulation present in real world climbing. What could additionally contribute to the underestimation of coarticulation in climbing is that the stPCs representations of the whole-body kinematics disregard the average speed of the climbing execution in the different climbing phases and the different trials (due to temporal normalization), a limitation mitigated by the fact that the overall velocity and accelerations profiles, as well as their relative values across all body joints, are preserved. Future works shall explore further how individual coarticulation strategies emerge when solving and executing more challenging climbing routes, by testing expert climbers alone.

A second critical limitation is the number of participants tested: given the large intra- and interindividual variability that characterize whole-body unconstrained motor behaviors, together with the need to group participants in three groups of expertise, a larger sample would be required to corroborate our findings. Third, assigning expertise levels to climbers is not obvious and the criteria we have adopted to segregate participants into three groups is to some extent arbitrary, particularly in those adopted to split climbers into *expert* and *beginners*, as the expertise level varies more plausibly on a continuous spectrum rather than in discrete clusters. Finally, this study leaves open the question of how coarticulation is realized at the computational and neural levels. As mentioned above, from a theoretical perspective, coarticulation might result from the activity of internal generative or forward models for motor control that are responsible for planning and steering sequences of movements (16,76,77). Future studies could more directly address the functional and neuronal underpinnings of the suggested internal models during sophisticated whole-body movements like climbing - which stands among the challenges to be faced in the context of future motor control research (2,10,78).

In conclusion, in this work we have provided novel evidence for coarticulation in the whole-body kinematics of participants conducting a climbing task, in which the sequence of arm movements was fully instructed. Coarticulation is observed independently of the level of expertise in the adjustment phase preparing the incoming limb moves but relates back to the previous move only for expert-climbers. These outcomes provide a new contribution to the current knowledge of the strict interplay between motor- and cognitive processes, specifically planning, and naturalistic motor behaviors.

## SUPPLEMENTAL MATERIAL

Supplementary material including the output of all statistical models and the figures of individual coarticulation maps from all participants are available at https://figshare.com/s/76adb56e6212ac928580.

## ACKNOWLEDGMENTS

We thank Laura Juppen and the team of intern students at the German Sport University Colone that supported AM with the lab setting, participant recruiting, and data collection.

## AUTHOR CONTRIBUTIONS

A.M., G.P. and A.dA. conceived and designed research, A.M. performed experiments, A.M. analyzed data, all authors interpreted results of experiments, A.M. prepared figures, A.M. drafted the manuscript, all authors edited and revised the manuscript, all authors approved final version of the manuscript.

